# Fine-tuning genomic models on rare splicing events identifies a novel biomarker of TDP-43 pathology

**DOI:** 10.64898/2026.10.07.757366

**Authors:** Cristina P. Martin-Linares, Aswathy Peethambaran Mallika, Parker S. Sandal, Tyler W. Martin, Meaghan Morris, Philip C. Wong, Jonathan P. Ling

**Affiliations:** Biomedical Engineering, Johns Hopkins University; Center for Computational Biology, Johns Hopkins University; Pathology, Johns Hopkins School of Medicine; Kavli Neuroscience Discovery Institute, Johns Hopkins University; Neuroscience, Johns Hopkins School of Medicine Baltimore, MD, USA

## Abstract

Genomic foundation models (GFMs) are pretrained on large genomic datasets, but their ability to predict rare disease-associated events after fine-tuning remains unclear. We tested this using TDP-43 cryptic exons, well characterized splicing events linked to neurodegenerative disease that are poorly represented in standard annotations. We compared the state-of-the-art GFM AlphaGenome with OpenSpliceAI, a specialized splice site prediction model. Fine-tuned OpenSpliceAI detected held-out cryptic exons in human and mouse and captured allele-specific splicing effects at the disease-associated *UNC13A* locus. In contrast, a splicing prediction head trained on AlphaGenome’s frozen pretrained backbone failed to predict cryptic exons. Unfreezing and fine-tuning all model weights, however, enabled AlphaGenome to generalize to held-out data. To investigate cryptic splicing in cell types that are difficult to study experimentally, we applied fine-tuned OpenSpliceAI across the genome to study cell type-specific genes. This led to the identification of new cryptic splicing events in *CLDN11* and *ARHGAP23*, which were subsequently validated in FTLD-TDP brain tissue. Notably, the cryptic exon in *CLDN11* is predicted to generate a neoepitope that could serve as an oligodendrocyte-specific biomarker of TDP-43 pathology. Our results show that even a state-of-the-art GFM can fail to generalize to rare splicing events, even with substantial pretraining. However, both foundation and specialized models that learn to generalize to rare biological events can have broad applications in biological discovery and medicine.

## 1 Introduction

Genomic foundation models (GFMs) are neural networks trained on large-scale genomics data to learn transferable representations of how DNA sequences map to biological function. Some GFMs use self-supervised training on genomic sequence alone [1–5]. Others use supervised training to predict multiple types of biological measurements from long genomic sequences [6–8]. Sequence-to-function models use large context windows to predict regulatory activity and variant effects that depend on distant interactions. Splicing is generally thought to depend primarily on local sequence context, and smaller specialized models perform well splice site prediction [9–14]. For AlphaGenome, performance on some benchmarks has been reported to exceed that of specialized splicing models [8].

Many genomic prediction tasks have limited training data and predicting pathogenic events that are rare or poorly annotated would require fine-tuning. Previous work suggests that GFMs can be successfully fine-tuned for new tasks beyond those represented in their original training data [15, 16]. However, fine-tuning a GFM would only be successful if the model learned features relevant to the new task during pretraining.

Indeed, strong cross-validation performance does not imply that GFMs will work well when applied to all biological settings. Recent benchmarks show that broad pretrained models do not always outperform specialized models, and in some cases provide only modest gains [17–19]. For personalized genomics, GFMs are better at predicting differences between genomic regions than differences between individuals, and often fail to predict the directionality of some variant effects [20, 21].

Alternative splicing prediction offers a useful setting to benchmark GFM performance because specialized models already perform well on standard splice site prediction, but often struggle on out-of-distribution events. Constitutive splice sites provide enough data to learn common splicing patterns. Models must also predict rare disease-associated changes that standard annotations do not capture.

We use TDP-43 cryptic exons to test whether fine-tuning can adapt pretrained models to events missing from standard annotations [22–27]. Cryptic exons are rare, absent from standard annotations, and only detected with specific experimental manipulations. We compared AlphaGenome (a GFM) and OpenSpliceAI (a specialized splicing model) across held-out human cryptic exons, mouse cryptic exons, and disease allele-specific predictions at the *UNC13A* genomic locus. We found that fine-tuning OpenSpliceAI leads to successful prediction of TDP-43 cryptic exons. In contrast, AlphaGenome does not predict cryptic exons when its backbone is frozen and only a splicing prediction head is fine-tuned. However, full model fine-tuning substantially improves AlphaGenome performance, approaching that of fine-tuned OpenSpliceAI. Full fine-tuning also appears to improve allele-specific predictions at the *UNC13A* locus.

Finally, we used the best performing model, fine-tuned OpenSpliceAI, and validated novel cryptic splicing events in *CLDN11* and *ARHGAP23* from FTLD-TDP brain tissue. The *CLDN11* event is predicted to produce a protein isoform with a new C-terminal sequence. This sequence is a candidate neoepitope for an oligodendrocyte-specific biomarker of TDP-43 pathology.

## 2 Benchmark and Fine-Tuning Setup

### 2.1 Cryptic exon benchmark and evaluation sets

We constructed a supervised fine-tuning benchmark from experimentally validated human TDP-43 cryptic exons [22–27]. These events are typically absent from standard splice annotations and become detectable after experimental perturbation. Each cryptic exon was represented by its genomic sequence context window and the experimentally determined positions of its 3’ splice site acceptor and 5’ splice site donor.

The benchmark includes three evaluation settings. First, we evaluate held-out human cryptic exons from the same supervised benchmark using gene level train, validation, and test partitions. Second, we test whether models fine-tuned on human cryptic exons can predict mouse TDP-43 cryptic splice sites. Third, we perform post-hoc human locus analyses at *UNC13A* and *HDGFL2*. At the ALS/FTD-associated *UNC13A* locus, we evaluate allele-specific predictions. At the *HDGFL2* locus, we evaluate cryptic splice site detection and false positive rates. Candidate genes and loci used for post-hoc evaluation were excluded from the fine-tuning splits.

### 2.2 Training setup

Training examples contained 15,000nt of one-hot encoded DNA sequence, with labels for the central 5,000nt. Labels specified per-base probabilities for background, acceptor, and donor channels. We added experimentally validated cryptic splice sites to the OpenSpliceAI labels, assigning 3’ splice sites to the acceptor channel and 5’ splice sites to the donor channel.

The full supervised dataset was partitioned into train, validation, and test sets using an 80/10/10 split at the gene level. All examples from one gene were assigned to the same partition. Validation examples were used for checkpoint selection, while held-out test examples were reserved for final evaluation. We repeated the analysis across independent gene-level splits. For each random partition, the human supervised benchmark contained 215 training genes, 26 validation genes, and 26 test genes, for a total of 267 fine-tuning and held-out benchmark genes. Four additional human genes were held out from all training datasets for downstream evaluation: HDGFL2, UNC13A, ACTL6B, and STMN2. In total, the human cryptic exon dataset used for fine-tuning and held-out benchmarking contained 411 annotated 3’ splice site acceptors and 406 annotated 5’ splice site donors [22, 24, 25, 27]. The mouse out-of-distribution benchmark contained 172 annotated 3’ splice site acceptors and 168 annotated 5’ splice site donors [23].

### 2.3 Model adaptation

We evaluated five model regimes: baseline OpenSpliceAI, fine-tuned OpenSpliceAI, baseline AlphaGenome, head-only fine-tuned AlphaGenome, and full-model fine-tuned AlphaGenome. Base-line models were evaluated without additional training. Fine-tuned OpenSpliceAI was initialized from the pretrained 10,000nt context model and updated on the cryptic exon training set using its three-channel splice site objective.

For AlphaGenome, we used a matched local-window adaptation setup. Each 15,000nt Open-SpliceAI window was converted to a 16,384nt AlphaGenome input while preserving the same central 5,000nt supervised interval. The converted AlphaGenome datasets reused the original labels without relabeling. We initialized AlphaGenome from its pretrained checkpoint and added a custom splice site head that predicts acceptor and donor probabilities at each base.

We considered two AlphaGenome fine-tuning protocols. In the head-only protocol, we froze the pretrained backbone and updated only the new splice site head. In the full-model protocol, we updated all AlphaGenome parameters and the splice site head. We compared these protocols to test whether cryptic splice site prediction required updates to the backbone.

This setup is intended as a matched local-transfer benchmark rather than an exhaustive evaluation of all AlphaGenome capabilities. In particular, it does not test AlphaGenome’s longer 1Mb context setting or all baseline splicing outputs. Sequence conversion, head architecture, and output conventions are described in Appendix B.

### 2.4 Evaluation metrics

Models produced base-resolution acceptor and donor probabilities across the supervised interval. For held-out and post-hoc cryptic exon analyses, we evaluated whether high-confidence predictions localized to experimentally validated cryptic splice site positions. Because cryptic splice sites are sparse relative to the number of possible intronic splice-like motifs, evaluation emphasized precision-recall behavior, average precision, and false positive burden rather than accuracy alone. For precision-recall analyses, validated cryptic splice sites were treated as positives, and the negative distribution was constructed from 5,000 randomly selected introns from GENCODE [28], corresponding to 9,965,479 evaluated negative base positions.

For mouse cryptic exon evaluation, we report precision-recall curves and average precision separately for 3’ splice site acceptor and 5’ splice site donor prediction. For the human *UNC13A* allele-specific analysis, we compare predictions across reference and alternate allele configurations and evaluate whether predicted cryptic splice site probabilities change relative to the healthy sequence. For the human *HDGFL2* locus analysis, we examine both cryptic exon-associated splice site recovery and the number of above-threshold predictions at background positions.

## 3 Experiments and Results

We evaluate the five model regimes described in Section 2 on mouse cryptic exons and at the human *UNC13A* and *HDGFL2* loci. We then use fine-tuned OpenSpliceAI to search for new cell type-specific cryptic splicing events and test the predictions in human brain tissue.

### 3.1 Fine-tuning improves mouse cryptic splice site prediction

We first evaluated whether models fine-tuned on human cryptic exons generalize to held-out mouse TDP-43 cryptic exons. For each model, we computed base-resolution acceptor and donor probabilities across the supervised interval and evaluated whether predictions localized to validated cryptic splice sites.

Figure 1 and Supplementary Table S1 show precision-recall performance for mouse cryptic exon splice site prediction. Fine-tuned OpenSpliceAI improves over baseline OpenSpliceAI for both 3’ splice site acceptor and 5’ splice site donor prediction. Baseline OpenSpliceAI achieves mean AP values of 0.019 *±* 0.005 for 3’ splice site prediction and 0.022 *±* 0.002 for 5’ splice site prediction, whereas fine-tuned OpenSpliceAI improves to 0.186 *±* 0.008 and 0.174 *±* 0.013, respectively. These improvements are observed across all five checkpoints.

**Figure 1.**
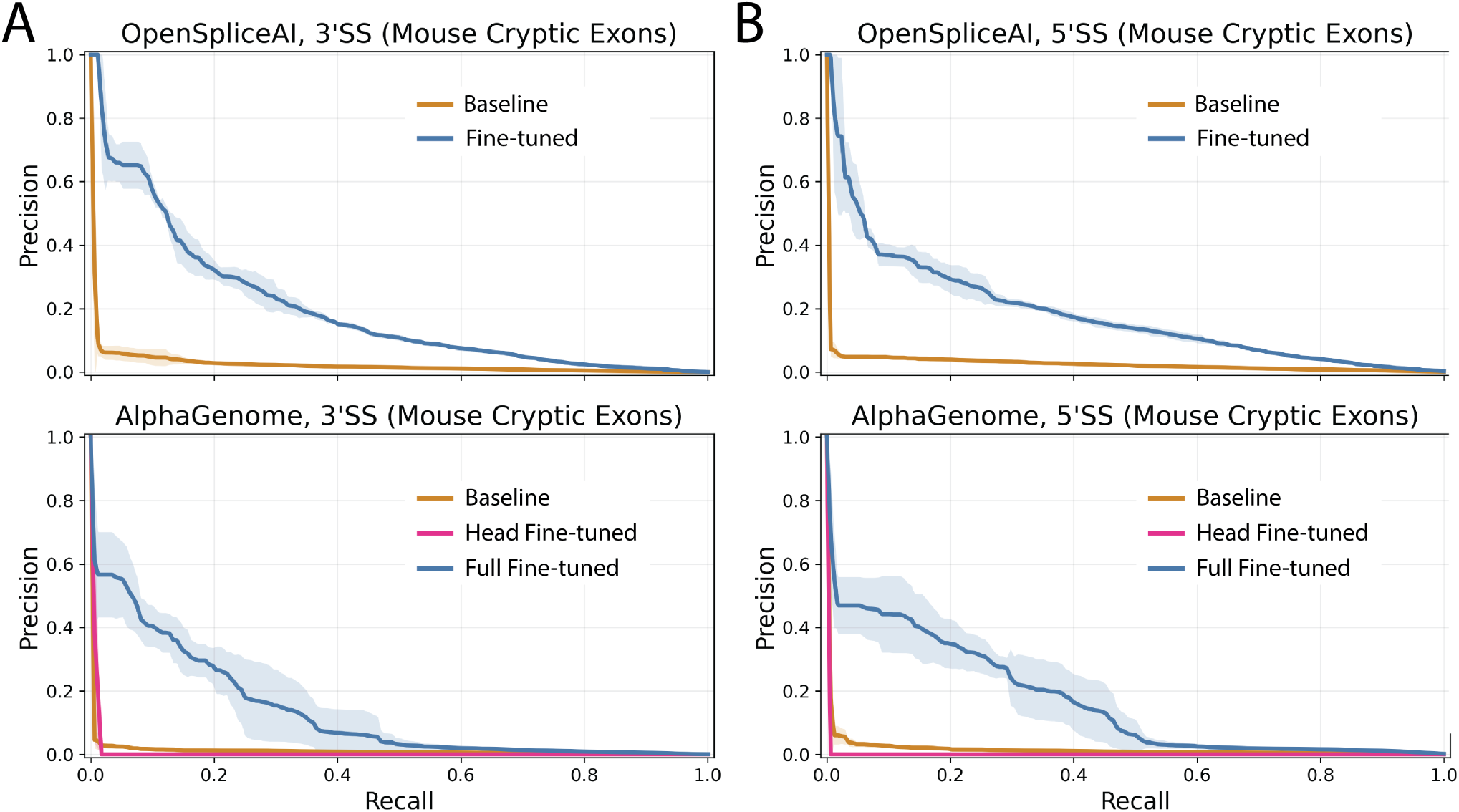
Fine-tuning improves cryptic splice site prediction of mouse TDP-43 cryptic exons. Precision-recall performance was evaluated for OpenSpliceAI and AlphaGenome on held-out mouse TDP-43 cryptic exon splice sites. (A) Precision-recall curves for 3’ splice site acceptor prediction. Fine-tuned OpenSpliceAI improves over baseline OpenSpliceAI, while only full-model fine-tuning improves AlphaGenome relative to baseline. Head-only fine-tuning of AlphaGenome fails to predict cryptic exons. (B) Precision-recall curves for 5’ splice site donor prediction show a similar pattern after fine-tuning.

AlphaGenome shows a different pattern. Baseline AlphaGenome performs weakly, with mean AP values of 0.008*±*0.001 for 3’ splice site prediction and 0.012*±*0.002 for 5’ splice site prediction. Head-only AlphaGenome fine-tuning fails to recover cryptic splice site signal, yielding AP values of 0.0 across all checkpoints for both splice site classes. In contrast, full-model AlphaGenome fine-tuning improves mean AP to 0.120 *±* 0.038 for 3’ splice site prediction and 0.147 *±* 0.031 for 5’ splice site prediction. Relative to baseline AlphaGenome, this corresponds to approximately 15.0*×* improvement for 3’ splice site prediction and 12.25*×* improvement for 5’ splice site prediction. Full-model AlphaGenome therefore recovers a substantial fraction of fine-tuned OpenSpliceAI performance, reaching approximately 65% of fine-tuned OpenSpliceAI mean AP for 3’ splice site prediction and 84% for 5’ splice site prediction. However, fine-tuned OpenSpliceAI remains stronger overall, especially for 3’ splice site prediction, where all full-model AlphaGenome checkpoints remain below all fine-tuned OpenSpliceAI checkpoints. For 5’ splice site prediction, full-model AlphaGenome is closer and the checkpoint ranges overlap, but its mean AP remains lower. In this setup, AlphaGenome required full-model fine-tuning to improve cryptic splice site prediction.

### 3.2 Allele-specific prediction at the *UNC13A* cryptic exon locus

We next tested whether fine-tuned models could detect cryptic splice sites at the ALS/FTD-associated *UNC13A* locus and predict allele-specific changes in splice site probability. We evaluated six genotypes: healthy, exon, intron, exon+intron, expansion, and all three alleles combined (exon+intron+expansion).

As shown in Figure 2, baseline OpenSpliceAI primarily assigns high splice site probability to constitutive splice sites and produces limited cryptic exon-associated signal. After fine-tuning, Open-SpliceAI predicts splice site signal in the cryptic exon region and shows allele-dependent changes relative to the healthy sequence.

**Figure 2.**
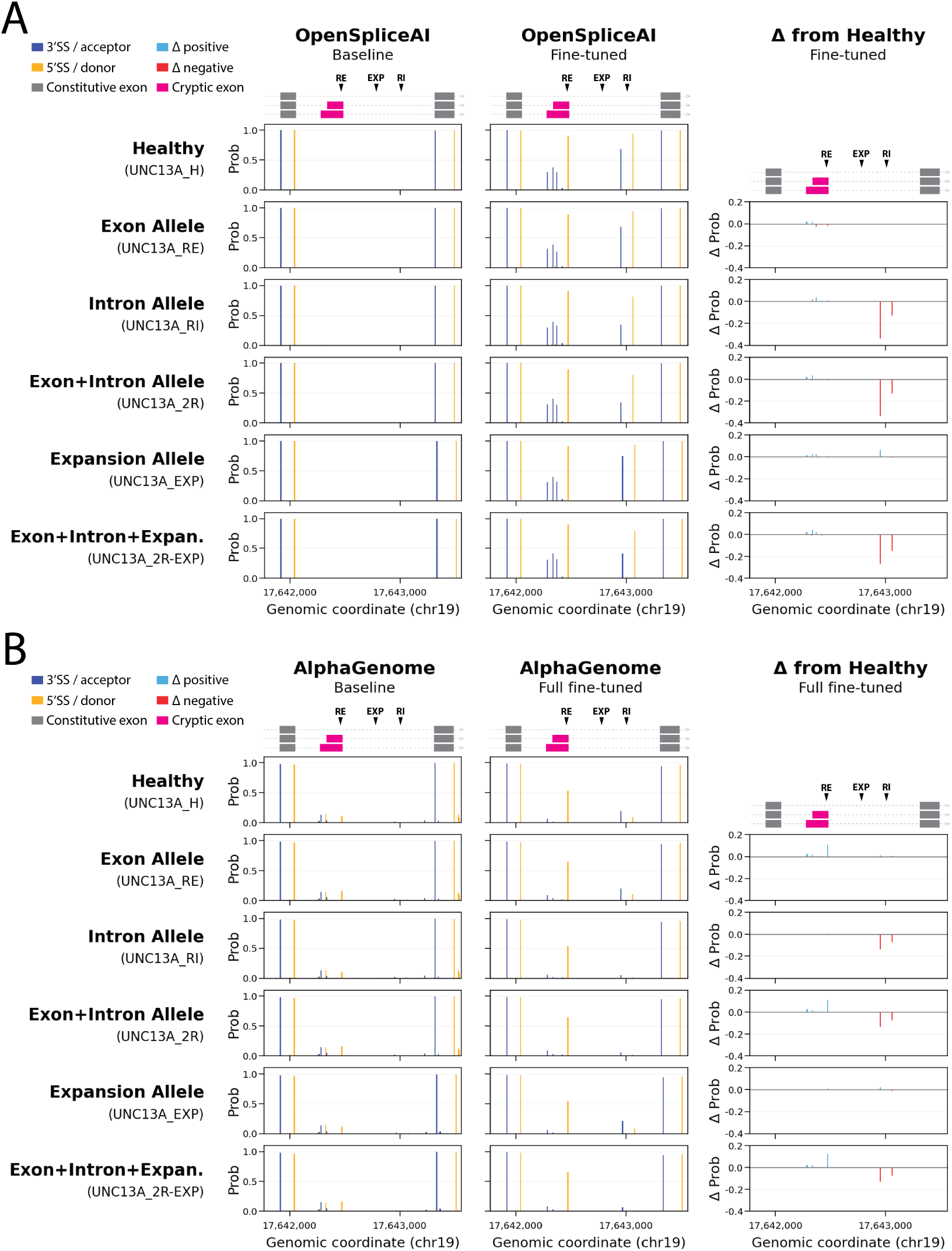
Allele-specific splice site predictions at the *UNC13A* cryptic exon locus. Certain genetic variants in the gene *UNC13A* are associated with neurodegenerative disease [24, 25]. Per-base splice site probabilities across the *UNC13A* cryptic exon locus are shown for six genotypes: healthy, exon, intron, exon+intron, expansion, and all three alleles. Gray box = constitutive exons, magenta box = cryptic exon. RE, EXP, and RI mark positions of the exon, expansion, and intron alleles, respectively. (A) Baseline and fine-tuned OpenSpliceAI predictions. The right column shows changes in fine-tuned predictions relative to the healthy sequence. (B) Baseline and full fine-tuned AlphaGenome predictions at the same locus, with the right column showing changes relative to the healthy sequence using the same Δ probability. While both fine-tuned models predict that RI containing alleles decrease neighboring splice site probabilities, only full-model fine-tuned AlphaGenome predicts an RE-associated increase in cryptic exon 5’ splice site probabilities. However, fine-tuned OpenSpliceAI better predicts the baseline cryptic splice site probabilities overall.

Full-model AlphaGenome fine-tuning also changes predictions at the *UNC13A* locus relative to the baseline model and produces signal near the cryptic exon region. However, this improvement is observed only when the AlphaGenome backbone is updated. The corresponding head-only AlphaGenome analyses, shown in the supplement, provide limited cryptic exon-associated signal.

### 3.3 Fine-tuning changes cryptic exon predictions and rates of false positives

Cryptic splice site prediction is highly imbalanced, with few validated positives and many plausible splice-like motifs in intronic sequence that are not used as cryptic splice sites. We therefore assessed recovery of validated cryptic splice sites alongside the number of above-threshold predictions at background positions.

Figure 3 shows per-base splice site prediction profiles across the *HDGFL2* locus. Baseline AlphaGenome produces broader intronic splice site signal across the evaluated region, consistent with a higher false positive burden. Head-only AlphaGenome fine-tuning reduces some diffuse background signal but does not reliably recover the cryptic exon-associated splice sites. Full-model AlphaGenome fine-tuning more strongly suppresses background predictions and produces clearer signal at the cryptic exon boundaries. OpenSpliceAI maintains a comparatively sparse prediction profile, and fine-tuning increases cryptic exon-associated signal without producing the same broad background activation observed for baseline AlphaGenome.

**Figure 3.**
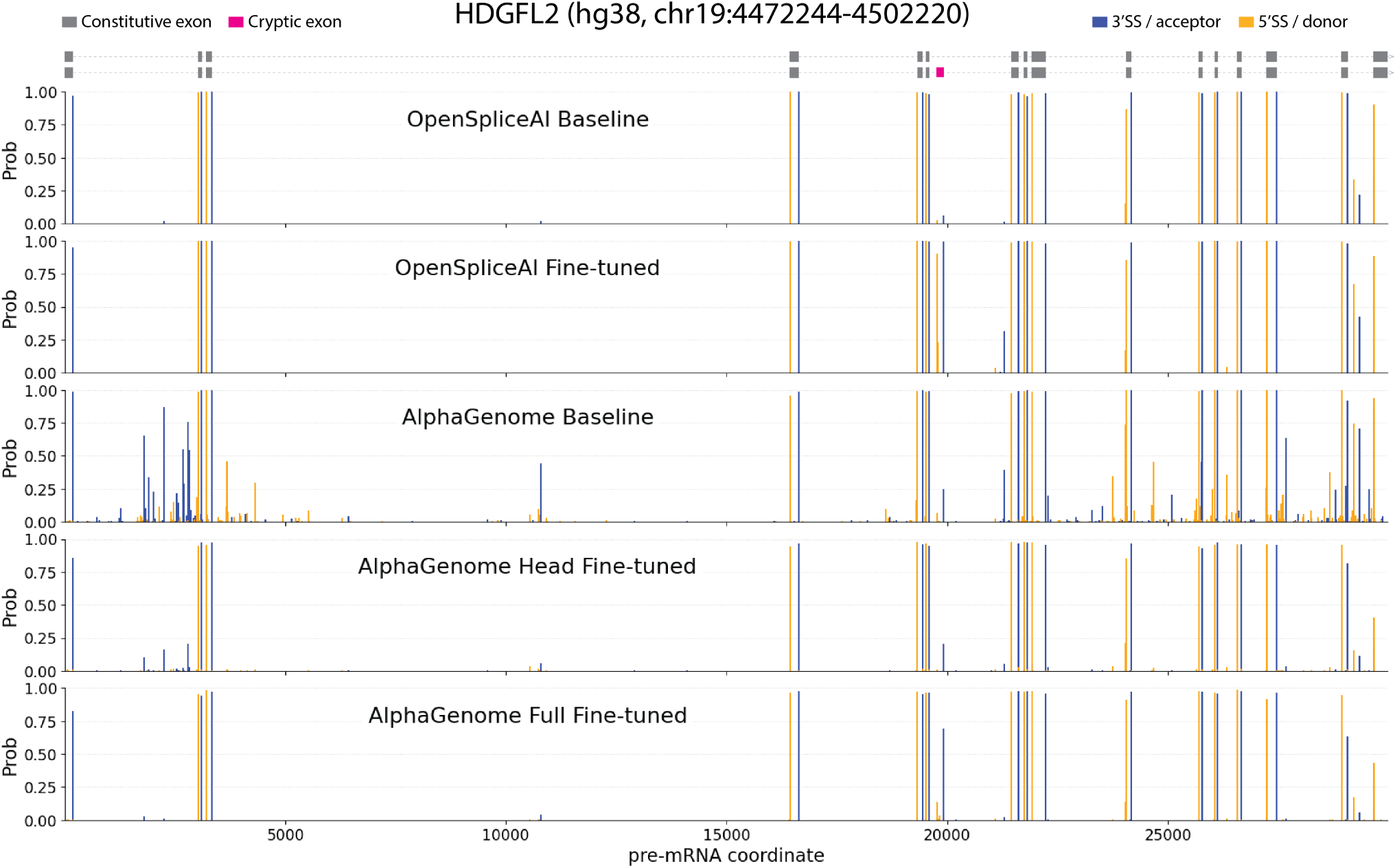
Splice-site prediction profiles across the *HDGFL2* locus. Per-base splice site probabilities are shown across the *HDGFL2* pre-mRNA region (hg38, chr19:4472244-4502220) for baseline and fine-tuned OpenSpliceAI and AlphaGenome models. Gray boxes mark constitutive exons and the magenta box marks the annotated *HDGFL2* cryptic exon. Blue vertical bars indicate predicted 3’ splice site acceptor probabilities, and orange vertical bars indicate predicted 5’ splice site donor probabilities. Each row shows model predictions from one inference regime: OpenSpliceAI baseline, OpenSpliceAI fine-tuned, AlphaGenome baseline, AlphaGenome head fine-tuned, and AlphaGenome full fine-tuned. Baseline AlphaGenome has the highest level of background predictions.

Figure 4 quantifies this effect across probability thresholds. Baseline AlphaGenome produces more above-threshold predictions at background positions, particularly at lower score thresholds. Fine-tuning reduces this burden, with the strongest improvement observed after full-model AlphaGenome fine-tuning. OpenSpliceAI shows lower background activation overall, both before and after fine-tuning.

**Figure 4.**
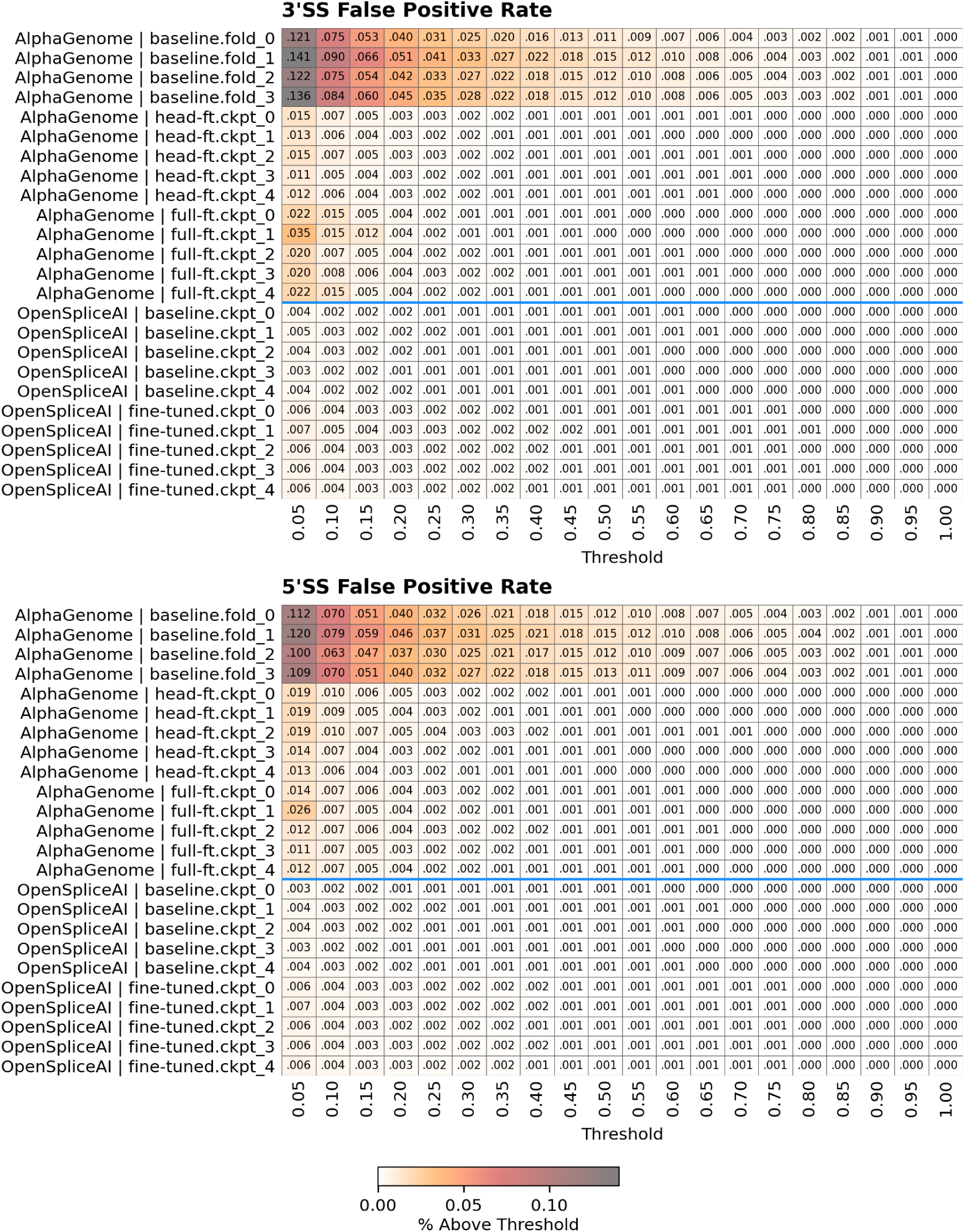
Fine-tuning reduces false positive splice site predictions across score thresholds. Heatmaps show the percentage of negative/background positions with predicted splice site probability above each threshold for 3’ splice site acceptors and 5’ splice site donors. Rows correspond to individual model folds or checkpoints for AlphaGenome and OpenSpliceAI, and columns correspond to probability thresholds from 0.05 to 1.00. Cell values report the percentage of evaluated positions exceeding each threshold, with darker shading indicating a higher false positive rate. Baseline AlphaGenome models produce more above-threshold predictions than fine-tuned AlphaGenome or OpenSpliceAI models, particularly at low thresholds. Blue horizontal lines separate AlphaGenome and OpenSpliceAI model groups.

Fine-tuned OpenSpliceAI and full-model fine-tuned AlphaGenome detect cryptic splice sites while limiting background predictions. Head-only AlphaGenome fine-tuning reduces background predictions but does not consistently detect the cryptic splice sites.

### 3.4 Genome-wide OpenSpliceAI predictions identify cryptic splicing in *CLDN11* and *ARHGAP23*

We used fine-tuned OpenSpliceAI to search for new cryptic splicing biomarkers in mature human brain cells. Some cell types, including oligodendrocytes, are difficult to model in vitro because current differentiation protocols generate cells that remain relatively immature. We ran baseline and fine-tuned OpenSpliceAI across all human genes and produced per-base acceptor and donor probabilities across all intronic sequences. We then selected splice sites that appeared higher predicted probabilities in the fine-tuned OpenSpliceAI model compared to baseline. We then cross-referenced these predictions with genes enriched in cell types that are difficult to model in vitro, which led to the identification of various cryptic exon candidates including those found in *CLDN11* and *ARHGAP23*.

To validate these candidates, we designed RT-PCR primers spanning the predicted cryptic splice junctions and tested postmortem brain tissue from FTLD-TDP and control cases and validated the cryptic exons in *CLDN11* and *ARHGAP23*. Primers targeting each event produced bands in middle frontal gyrus (MFG) samples from a subset of FTLD-TDP cases, but not from control cases (Figure 5). We detected no bands in cerebellar tissue from either group. Detection varied among FTLD-TDP cases and correlated with semiquantitative grading of oligodendroglial TDP-43 pathology, suggesting that these events may reflect cell type-specific pathology and splicing changes. As positive controls, primers targeting known TDP-43 cryptic exons in *HDGFL2*, *UNC13A*, and *STMN2* produced the expected bands only in FTLD-TDP samples, confirming the presence of TDP-43 pathology in the tissue used for this analysis.

**Figure 5.**
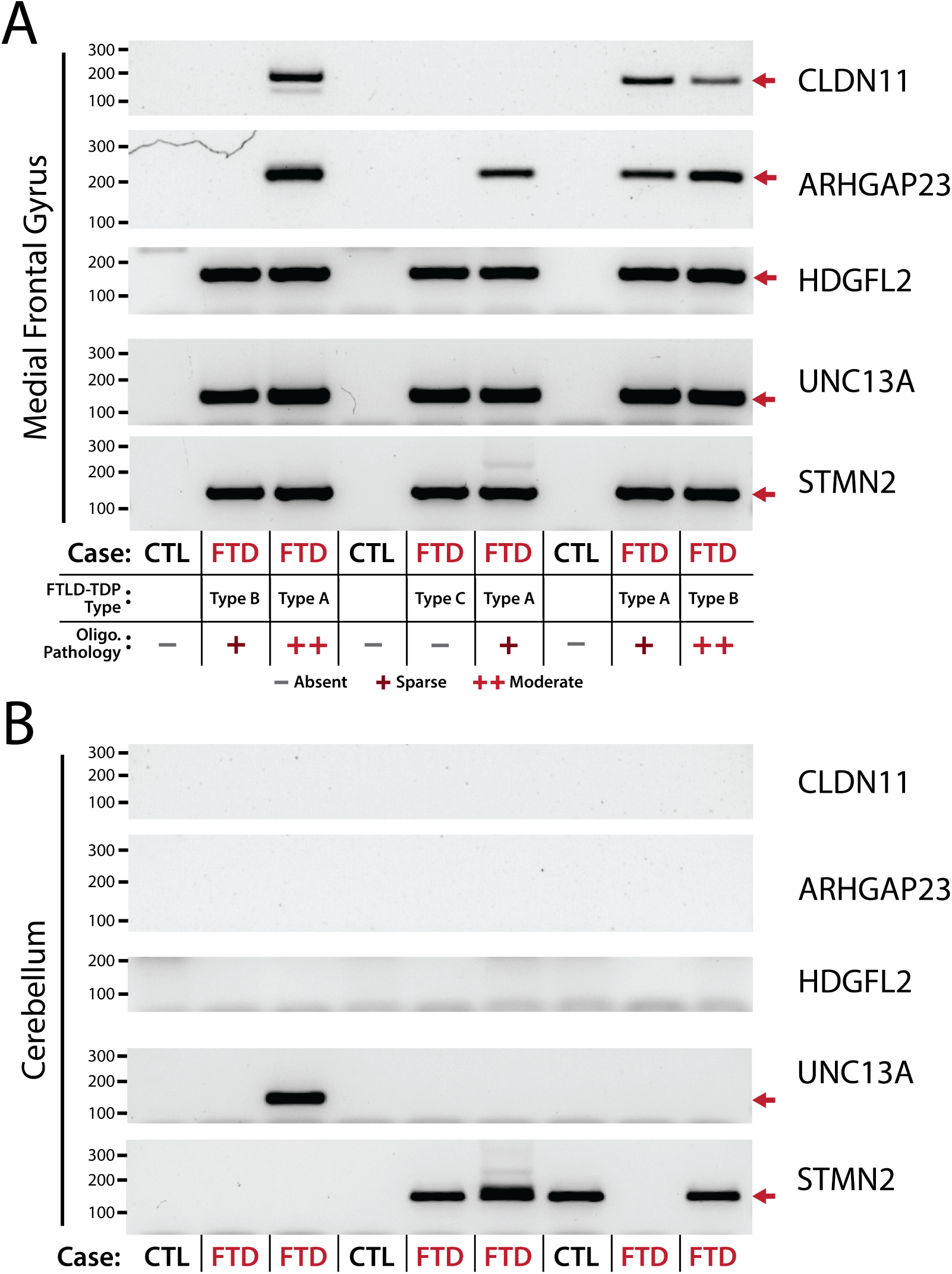
Human brain validation of the *CLDN11* and *ARHGAP23* cryptic splicing events discovered using fine-tuned OpenSpliceAI. RT-PCR was performed using primers designed to amplify single band cryptic splicing amplicons for *CLDN11*, *ARHGAP23*, and positive control TDP-43 cryptic exons for *HDGFL2*, *UNC13A*, and *STMN2*. Red arrows indicate the expected size for the cryptic RT-PCR amplicons. (A) Middle frontal gyrus samples from control (CTL) and FTLD-TDP (FTD) cases. FTLD-TDP subtype and semi-quantitative oligodendroglial TDP-43 pathology are indicated below the lanes (*−* = absent, + = sparse, ++ = moderate). *CLDN11* and *ARHGAP23* cryptic amplicons are detected in a subset of FTLD-TDP cases and are absent from controls, whereas positive control cryptic amplicons are detected across all FTLD-TDP samples. (B) Cerebellum samples from the corresponding case series, *CLDN11* and *ARHGAP23* cryptic amplicons are not detected.

Sanger sequencing of gel excised RT-PCR products confirmed that the amplified bands corresponded to the predicted *CLDN11* and *ARHGAP23* cryptic splice junctions, rather than nonspecific amplicons (Supplementary Figure S9). The predicted *CLDN11* event uses an alternative 3’ splice site in the final intron. This event is predicted to generate a protein-coding transcript that replaces the canonical C-terminal coding sequence with a novel C-terminal region. Boltz-2 protein structure predictions suggest that the cryptic *CLDN11* isoform retains a folded N-terminal structure resembling wildtype *CLDN11*, but introduces a unique C-terminal domain absent from the canonical protein (Supplementary Figure S10).

The predicted C-terminal region is a candidate neoepitope for antibodies that distinguish the cryptic isoform from wild-type CLDN11. In the brain, CLDN11 is highly specific to mature oligodendrocytes and myelin. The predicted cryptic protein is therefore a candidate for fluid biomarker studies of oligodendrocyte-specific TDP-43 pathology.

## 4 Conclusion

In this benchmark, a new prediction head on the frozen AlphaGenome backbone did not recover TDP-43 cryptic splice sites. Updating all AlphaGenome weights improved prediction, including allele-specific effects at the *UNC13A* locus. Fine-tuned OpenSpliceAI performed better overall on direct cryptic splice site prediction. Full-model fine-tuned AlphaGenome may be useful for allele-specific analyses. We used fine-tuned OpenSpliceAI to identify a new cryptic splicing event in *CLDN11* and validated its splice junction in FTLD-TDP brain tissue. The event is predicted to produce a new C-terminal sequence. This sequence is a candidate neoepitope for an oligodendrocyte-specific biomarker of TDP-43 pathology. The benchmark can test how well pretrained models adapt to rare splicing events. Whether including these events during pretraining improves prediction remains an open question.

## 5 Acknowledgements

This work was supported in part by the NIH grants (UG3/UH3NS115608, R33NS115161 and R01NS095969 to P.C.W.), and the Department of Defense (AL230138 to P.C.W.). This study was also supported by the Johns Hopkins Alzheimer’s Disease Research Center (P30AG066507).

## Supplementary Information

### A Additional results

#### Precision analyses

Supplementary Table S1. records mean AP for the mouse benchmark. Individual AP values and precision-recall curves are shown in Supplementary Figures S1, S2, and S3.

#### *UNC13A* cryptic exon predictions across models

Supplementary Figures S4 and S5 show individual model predictions for the allele-specific analysis in Figure 2.

#### *HDGFL2* cryptic exon predictions across models

Supplementary Figures S6, S7, and S8 show individual model predictions underlying Figure 3.

#### CLDN11 and ARHGAP23 validation

### B Supplementary methods

#### Data partitions

We split genes before generating sequence windows and used the same partitions for both models. The downstream evaluation genes listed in the main text were excluded before splitting. We verified that the gene lists did not overlap between partitions.

#### Sequence conversion

We converted 15,000nt OpenSpliceAI windows to 16,384nt AlphaGenome inputs while preserving the central 5,000nt labels. We reconstructed source sequences using exact 10,000nt overlaps between adjacent windows. Conversion stopped if the overlaps did not match. Each supervised interval was centered in its new input window, with zero-padding only beyond sequence boundaries.

#### AlphaGenome splice head

At each base, the head applies a 256-unit hidden layer with a ReLU activation and a two-unit output layer. The same head is used for head-only and full-model fine-tuning.

#### Inference and output channels

Baseline AlphaGenome uses its native SPLICE_SITES output, which has separate acceptor and donor tracks for each strand [8]. We selected the tracks for the inference strand and cropped them to the central 5,000nt interval. Fine-tuned AlphaGenome uses unstranded acceptor and donor outputs. We trained these outputs on the corresponding OpenSpliceAI labels after discarding the background channel. Low probabilities in both channels represent background. All models were compared using acceptor and donor probabilities over the same central 5,000nt interval.

#### Code and computing resources

All experiments were conducted on NVIDIA RTX PRO 6000 GPUs, with each fine-tuning run completing in under one hour. The training splits, held-out datasets, and scripts used for training and inference are available at https://github.com/cristinamartinlinares/gfm-splicing-benchmark.

---

**Figure S1.**
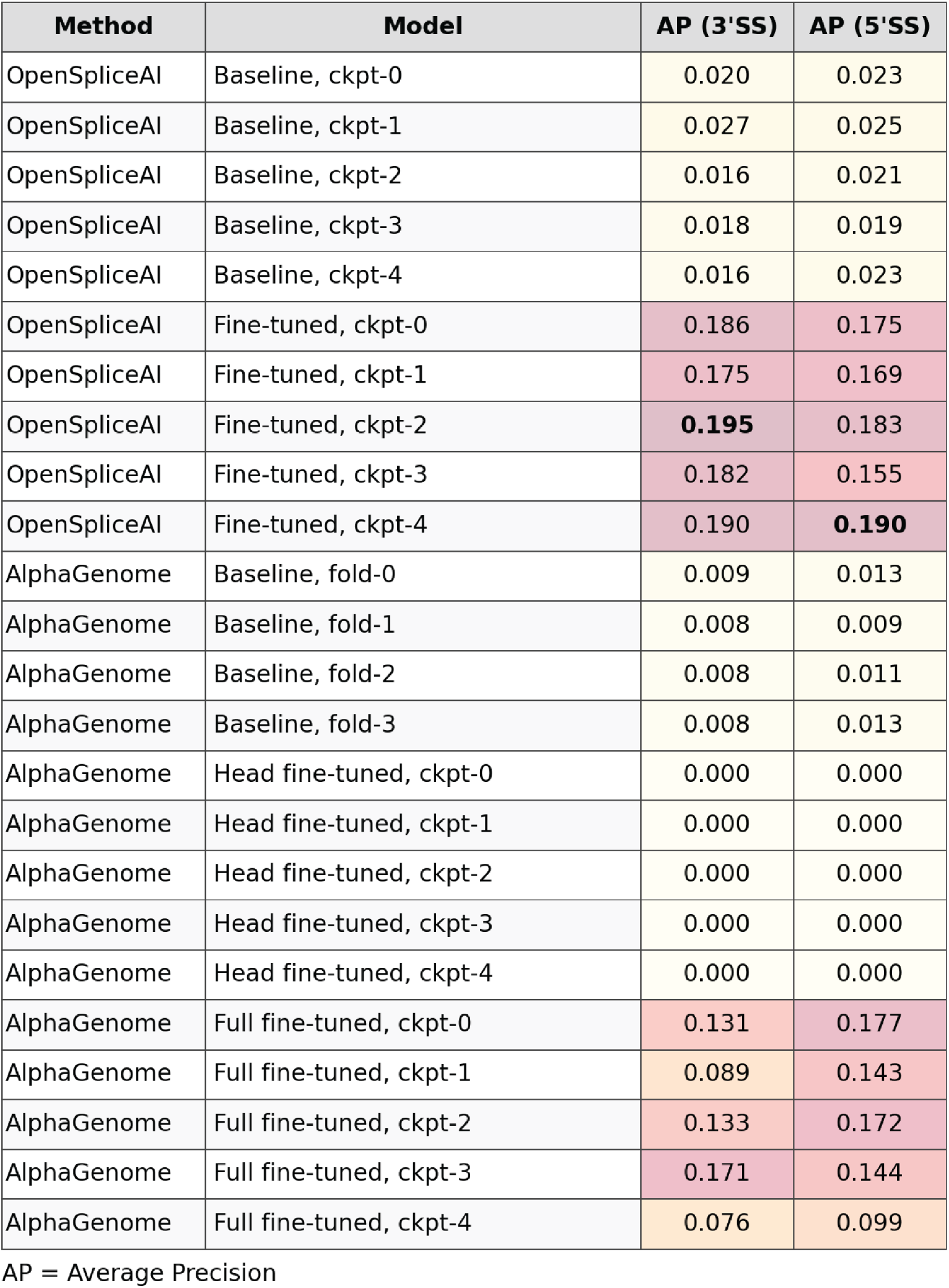
Average precision for mouse cryptic splice site prediction. Table summarizing average precision (AP) for 3’ splice site acceptor and 5’ splice site donor prediction across model folds or checkpoints. Precision–recall curves are shown in Figure 1; mean AP values are reported in Supplementary Table S1.

**Table S1.** Average precision on mouse TDP-43 cryptic exon splice site prediction. Values report mean *±* standard deviation across evaluated folds or checkpoints. Baseline AlphaGenome is evaluated across four folds; OpenSpliceAI baseline, OpenSpliceAI fine-tuned, and AlphaGenome fine-tuned regimes are evaluated across five checkpoints. Individual checkpoint AP values are reported in Supplementary Figure S1.

| Method | Regime | AP 3' splice site / acceptor | AP 5' splice site / donor |
| --- | --- | --- | --- |
| OpenSpliceAI | Baseline | 0.019 ± 0.005 | 0.022 ± 0.002 |
| OpenSpliceAI | Fine-tuned | **0.186 ± 0.008** | **0.174 ± 0.013** |
| AlphaGenome | Baseline | 0.008 ± 0.001 | 0.012 ± 0.002 |
| AlphaGenome | Head-only fine-tuned | 0.000 ± 0.000 | 0.000 ± 0.000 |
| AlphaGenome | Full-model fine-tuned | 0.120 ± 0.038 | 0.147 ± 0.031 |

**Figure S2.**
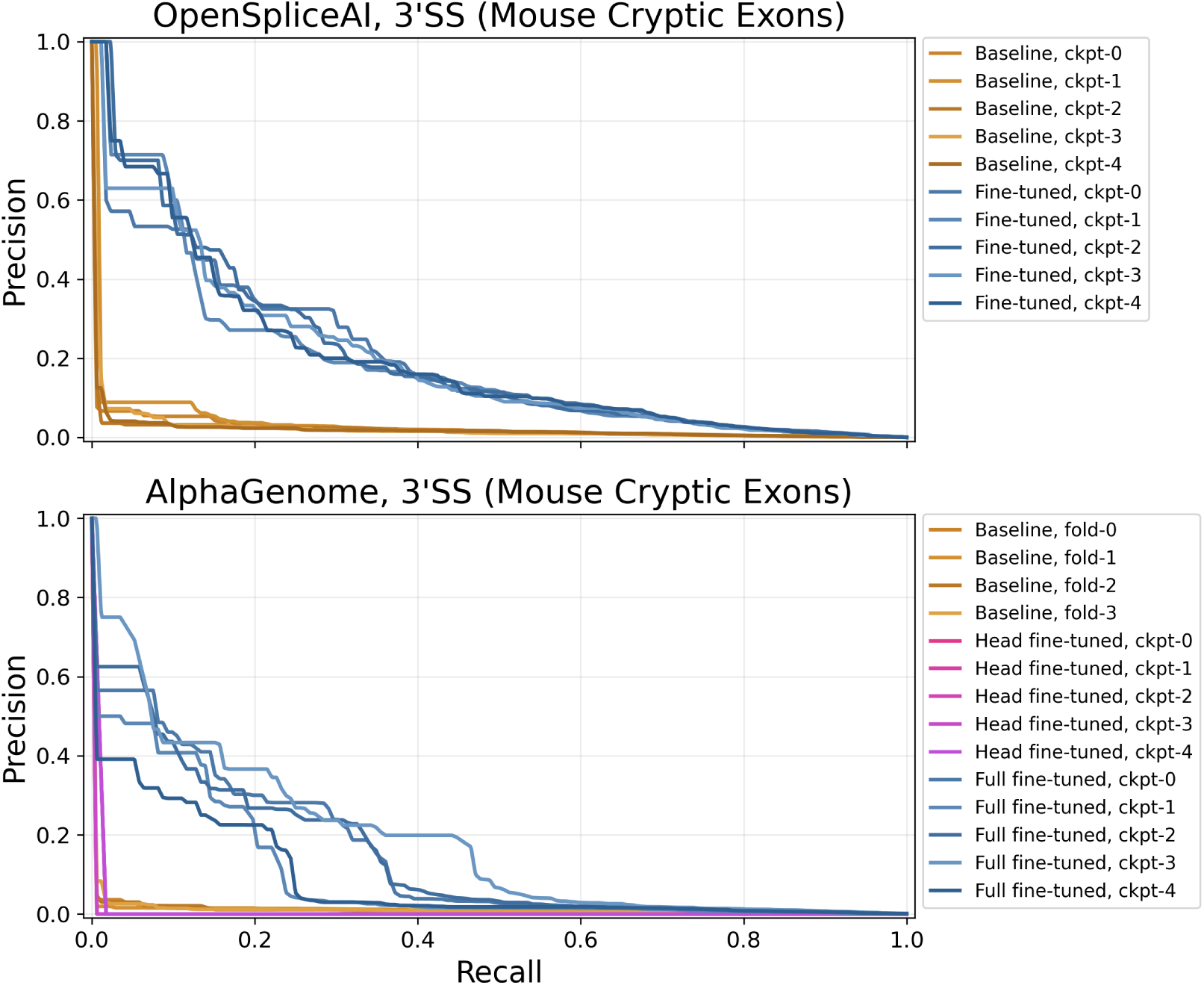
Individual 3’ splice site precision-recall curves underlying Figure 1, shown separately for each model fold or checkpoint.

**Figure S3.**
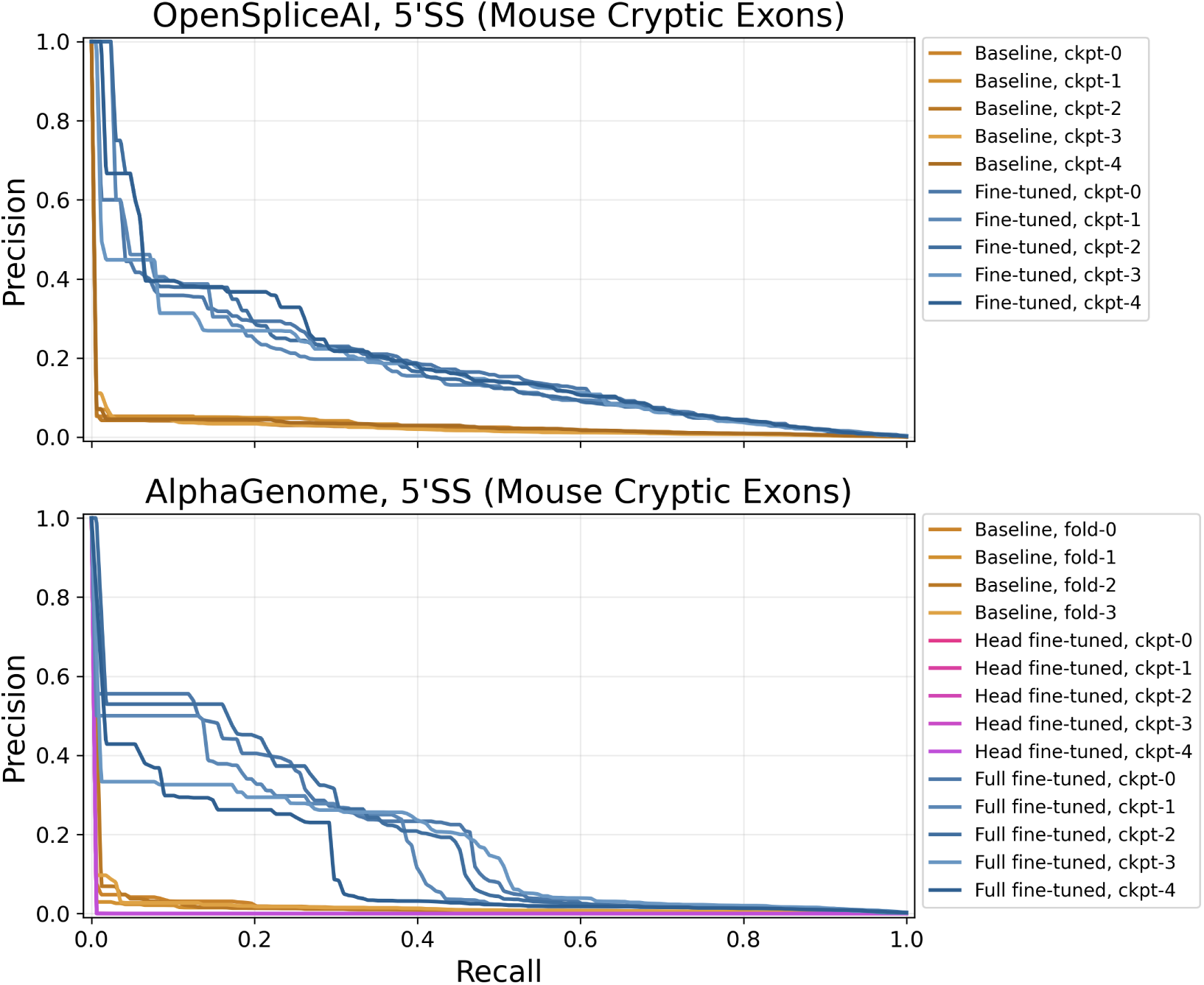
Individual 5’ splice site precision-recall curves underlying Figure 1, shown separately for each model fold or checkpoint.

**Figure S4.**
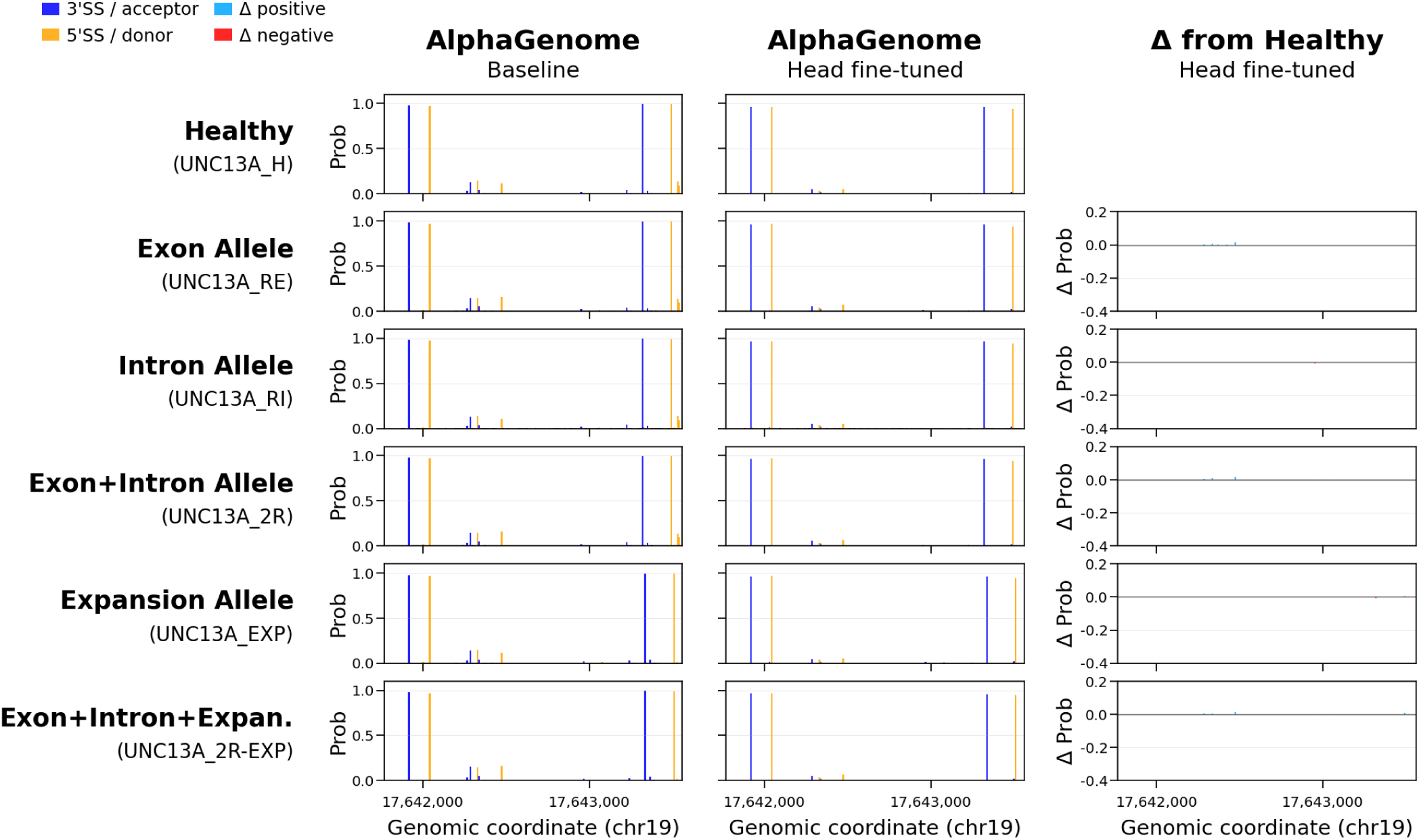
Head-only fine-tuning of AlphaGenome at the *UNC13A* locus. Head-only fine-tuned AlphaGenome predicts no cryptic exon splice sites or allele-specific splicing changes. Compare these predictions with Figure 2.

**Figure S5.**
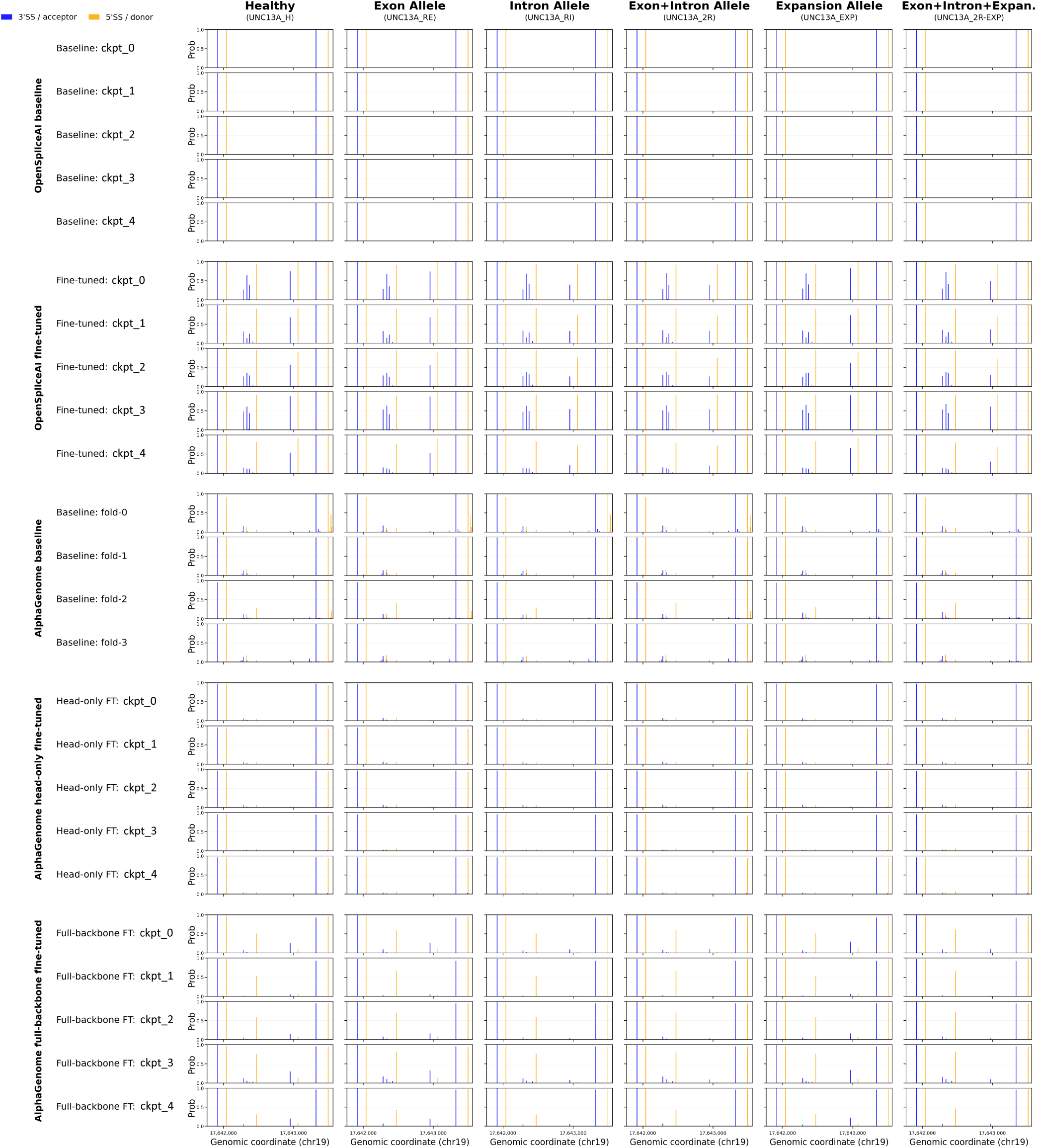
Individual model predictions at the *UNC13A* locus. Same allele-specific analysis as Figure 2, shown separately for each model fold or checkpoint.

**Figure S6.**
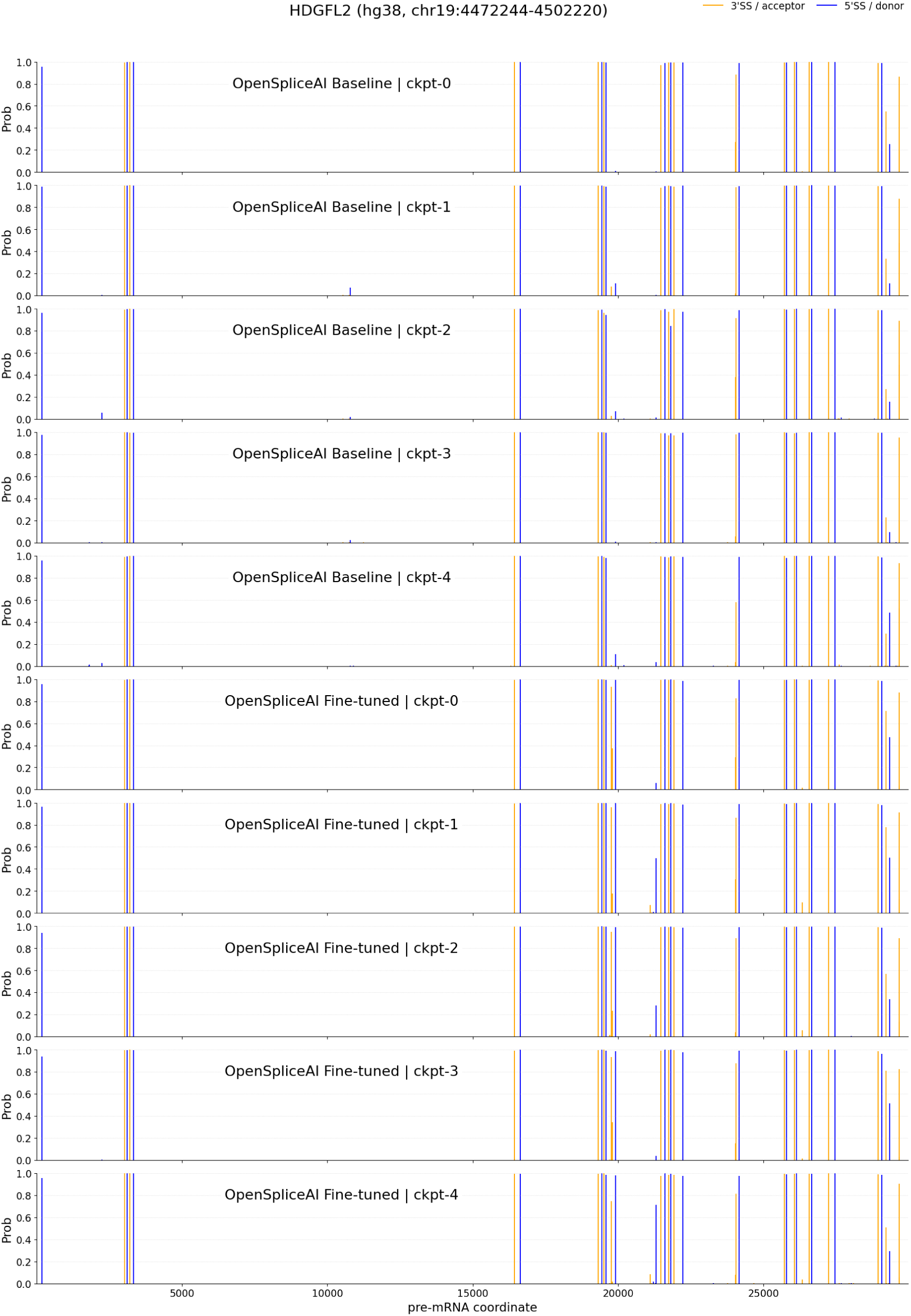
OpenSpliceAI predictions at the *HDGFL2* locus. Individual OpenSpliceAI baseline and OpenSpliceAI fine-tuned model predictions underlying Figure 3.

**Figure S7.**
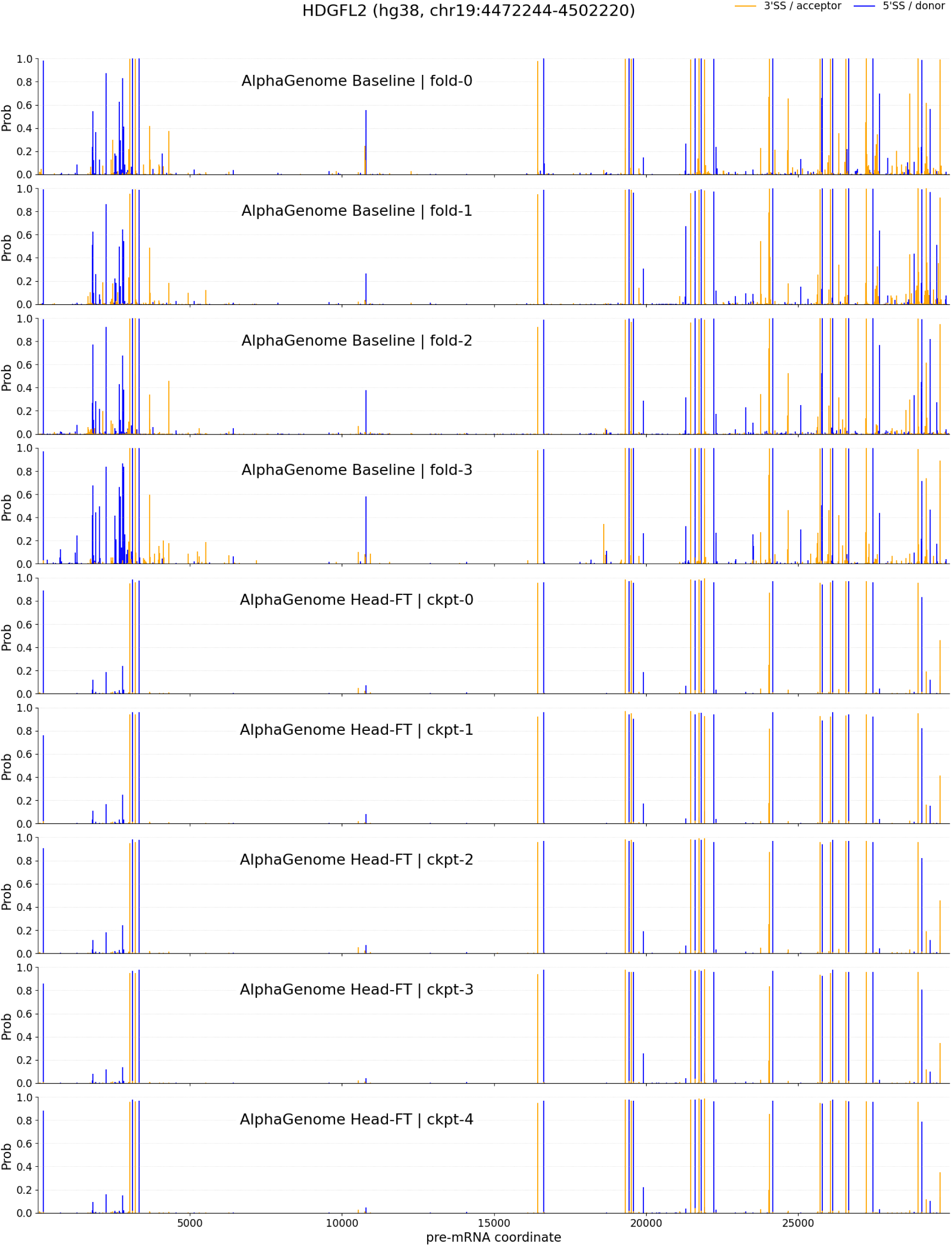
AlphaGenome head-only fine-tuning at the *HDGFL2* locus. Individual AlphaGenome baseline and AlphaGenome head-only fine-tuned model predictions underlying Figure 3.

**Figure S8.**
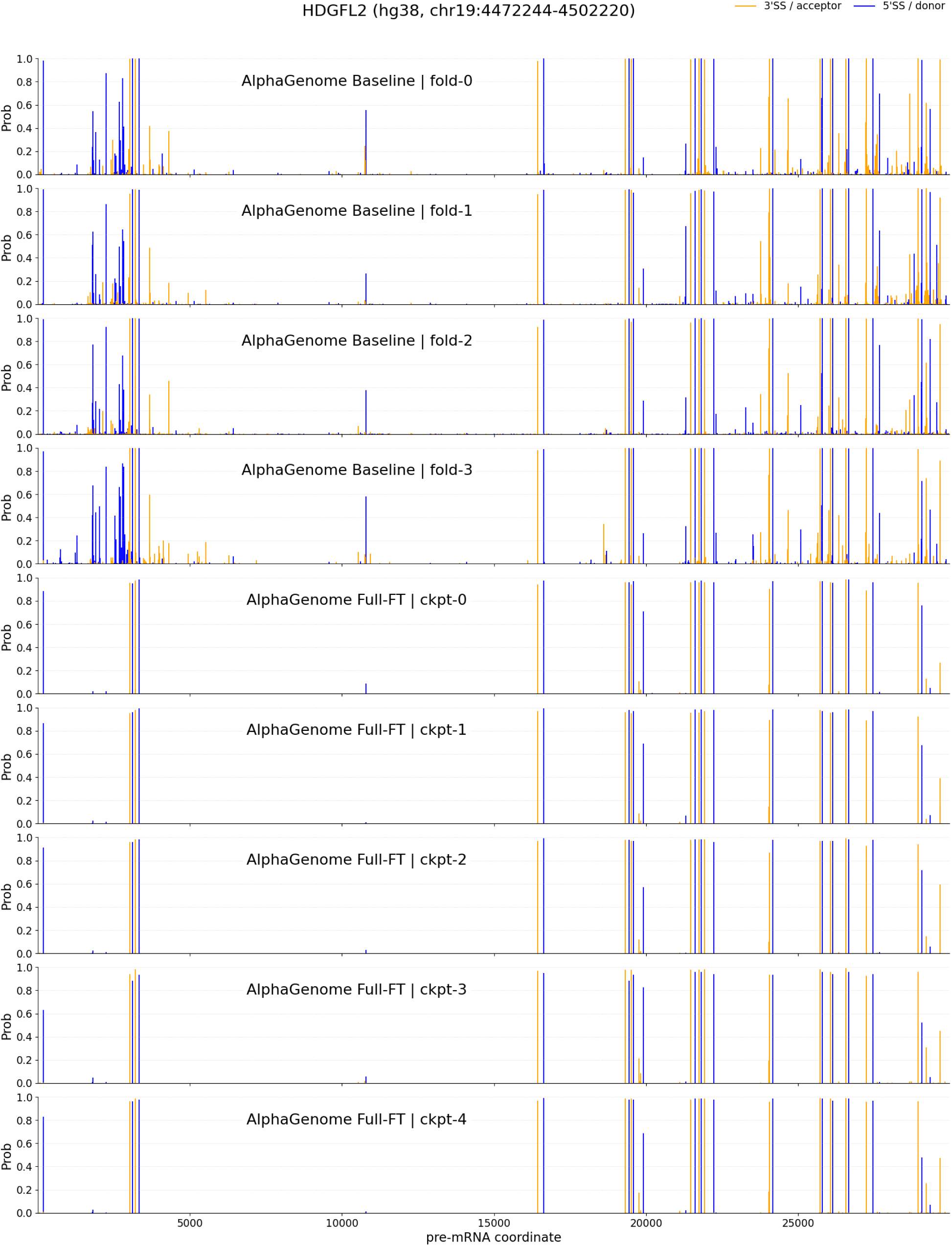
AlphaGenome full-model fine-tuning at the *HDGFL2* locus. Individual AlphaGenome baseline and AlphaGenome full-model fine-tuned predictions underlying Figure 3.

**Figure S9.**
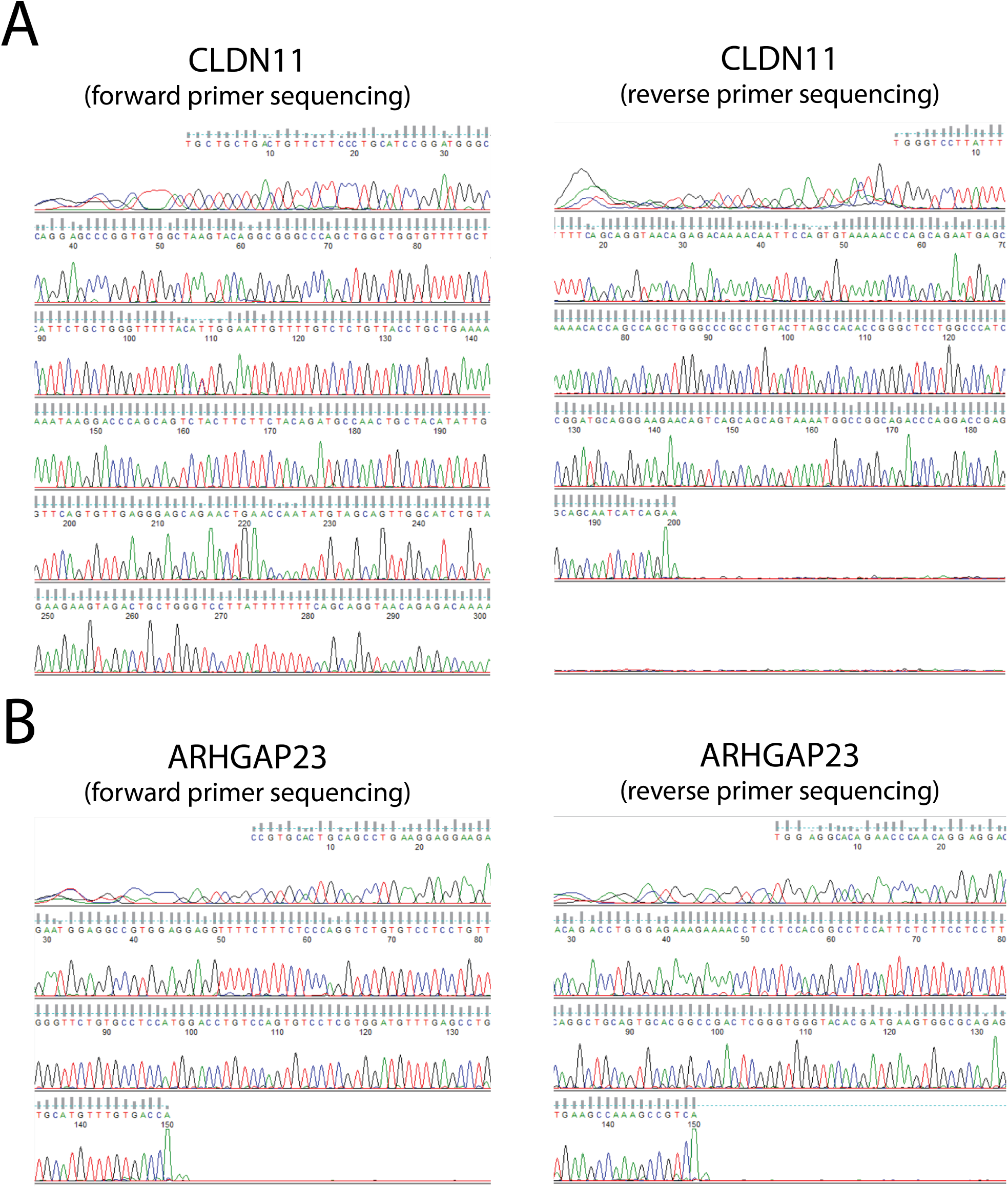
Sanger sequencing confirmation of *CLDN11* and *ARHGAP23* RT-PCR amplicons. Gel excised RT-PCR bands from Figure 5 were sequenced using forward and reverse primers. Chromatograms confirm that the predicted *CLDN11* (A) and *ARHGAP23* cryptic splicing amplicons match the predicted cryptic splice junctions.

**Figure S10.**
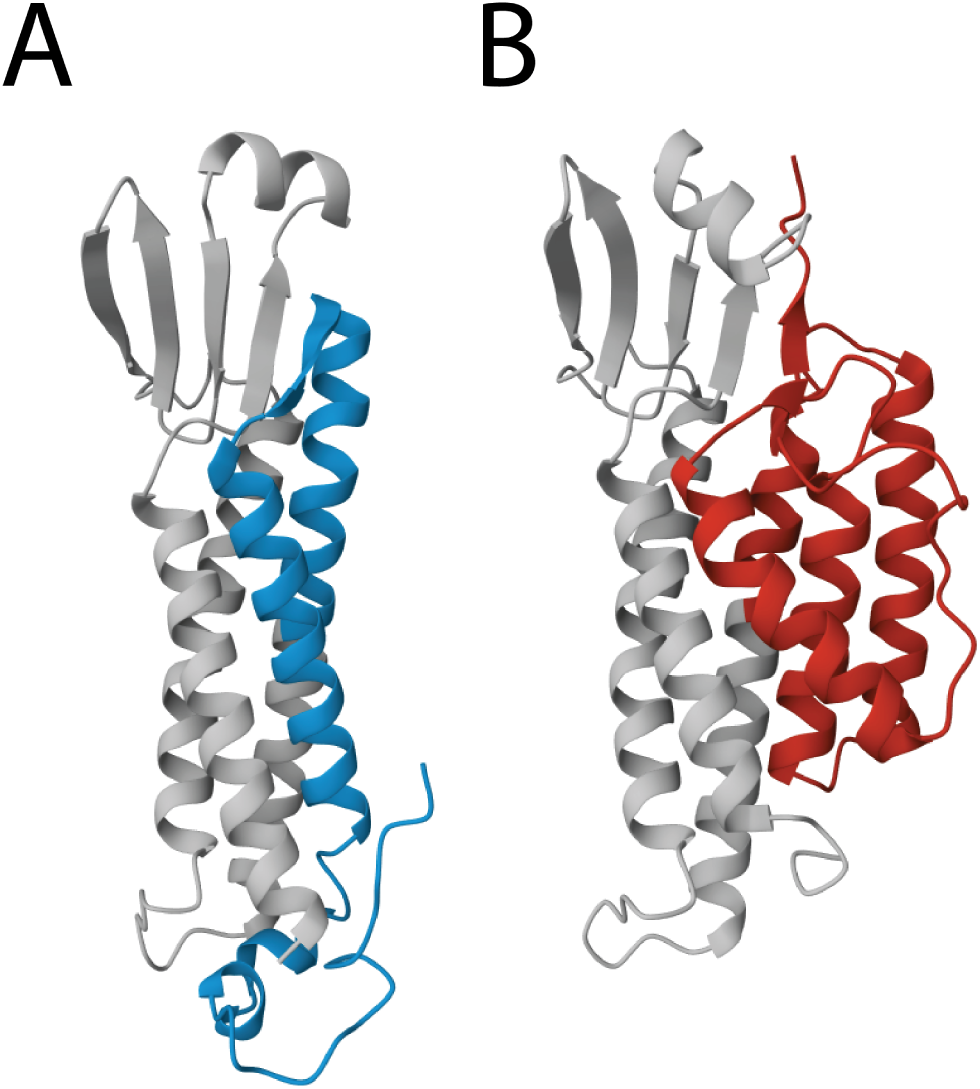
Predicted structural consequence of the cryptic alternative 3’ splice site in *CLDN11*. Boltz-2 protein structure predictions are shown for wildtype CLDN11 (A) and the cryptic CLDN11 isoform (B) generated by the alternative 3’ splice site in the final intron. Gray highlights indicate shared sequences, blue indicates sequences unique to wildtype *CLDN11*, and red indicates sequences unique to cryptic *CLDN11*.

## Notes

https://github.com/cristinamartinlinares/gfm-splicing-benchmark

### Competing Interest Statement

The authors have declared no competing interest.

